# Prefusion spike protein stabilization through computational mutagenesis

**DOI:** 10.1101/2020.06.17.157081

**Authors:** Dong Yan Zhang, Jian Wang, Nikolay V. Dokholyan

**Author notes:** Corresponding author: Nikolay V. Dokholyan,.

## Abstract

A novel severe acute respiratory syndrome (SARS)-like coronavirus (SARS-CoV-2) has emerged as a human pathogen, causing global pandemic and resulting in over 400,000 deaths worldwide. The surface spike protein of SARS-CoV-2 mediates the process of coronavirus entry into human cells by binding angiotensin-converting enzyme 2 (ACE2). Due to the critical role in viral-host interaction and the exposure of spike protein, it has been a focus of most vaccines’ developments. However, the structural and biochemical studies of the spike protein are challenging because it is thermodynamically metastable^1^. Here, we develop a new pipeline that automatically identifies mutants that thermodynamically stabilize the spike protein. Our pipeline integrates bioinformatics analysis of conserved residues, motion dynamics from molecular dynamics simulations, and other structural analysis to identify residues that significantly contribute to the thermodynamic stability of the spike protein. We then utilize our previously developed protein design tool, Eris, to predict thermodynamically stabilizing mutations in proteins. We validate the ability of our pipeline to identify protein stabilization mutants through known prefusion spike protein mutants. We finally utilize the pipeline to identify new prefusion spike protein stabilization mutants.

## INTRODUCTION

The ongoing outbreak of the novel coronavirus^2–4^, which causes fever, severe respiratory illness, and pneumonia, poses a major public health and governance challenges. The emerging pathogen has been characterized as a new member of the betacoronavirus genus (SARS-CoV-2)^5,6^, closely related to several bat coronaviruses and to severe acute respiratory syndrome coronavirus (SARS-CoV)^7,8^. Compared with SARS-COV, SARS-CoV-2 appears to be more readily transmitted from human to human, spreading to multiple continents and leading to the World Health Organization (WHO)’s declaration of a pandemic on March 11, 2020^9^. According to the WHO, as of June 9, 2020, there had been >7,000,000 confirmed cases globally, leading to >400,000 deaths.

In the initial step of the infection, the coronavirus enters the host cell by binding to a cellular receptor and fusing the viral membrane with the target cell membrane^10,11^. For SARS-CoV-2, this process is mediated by spike glycoprotein which is a homo-trimer^12^. In the process of viruses fusing to host cells, the spike protein undergoes structural rearrangement and transits from a metastable prefusion conformational state to a highly stable post-fusion conformational state^13,14^. The spike protein comprises of two functional units, S1 and S2 subunits; when fused to the host cell, the two subunits are cleaved. The S1 subunit is responsible for binding to the angiotensin-converting enzyme 2 (ACE2)^15–17^ receptor on the host cell membrane and it contains the N-terminal domain (NTD), the receptor-binding domain (RBD) and the C-terminal domain (CTD). NTD in the S1 subunit assists recognize sugar receptors. RBD in the S1 subunit is critical for the binding of coronavirus to the ACE-2 receptor^18–21^. CTD in the S1 subunit could recognize other receptors^22^. The binding of RBD to ACE2 facilitates the cleavage of the spike protein and promotes the dissociation of the S1 subunit from the S2 subunit^23^. S2 contains two heptad repeats (HR1 and HR2), a fusion peptide, and a protease cleavage site (S2’). The dissociation of S1 induces S2 to undergo a dramatic structural change to fuse the host and viral membranes. Thus, the spike protein serves as a target for development of antibodies, entry inhibitors and vaccines^24^. Coronavirus transits from a metastable prefusion state to a highly stable post-fusion state as part of the spike protein’s role in membrane fusion. The instability of the prefusion state presents a significant challenge for the production of protein antigens for antigenic presentation of the prefusion antibody epitopes that are most likely to lead to neutralizing responses. Thus, since the prefusion spike protein exists in a thermodynamically metastable state^1^, a stabilized mutant conformation is critical for the development of vaccines and drugs.

Computational mutagenesis is an effective approach to finding mutations that are able to stabilize proteins. We have previously developed a protein design platform, Eris^25,26^, which utilizes a physical force field^27^ for modeling inter-atomic interactions, as well as fast side-chain packing and backbone relaxation algorithms to enable efficient and transferrable protein molecular design. Originally, Eris has been validated on 595 mutants from five proteins, corroborating the unbiased force field, side-chain packing and backbone relaxation algorithms. In many later studies, Eris has been validated through prediction of thermodynamically stabilizing or destabilizing mutations^28–33^, and direct protein design efforts^34–37^.

In this work, we propose a pipeline to automatically stabilize spike proteins through computational mutagenesis. Within the pipeline, we first analyze the conservation score and solvent accessible surface area (SASA) of residues in the protein. We then perform discrete molecular dynamics (DMD)^38–41^ simulations to calculate the root mean square fluctuation (RMSF) of residues to analyze their flexibility. Based on this information, we select appropriate residues (2 < Conservation Score < 5; SASA > 0.4; RMSF > 3.5 Å; see Discussion) as mutation sites. We subject the selected residues to computational redesign using Eris to find the stabilizing mutations by calculating the change in free energy ΔΔ*G* = Δ*G*_*mut*_ *−* Δ*G*_*WT*_, where Δ*G*_*mut*_ and Δ*G*_*WT*_ are the free energies of the mutant protein and wild type proteins correspondingly. We utilize this pipeline to identify stabilization mutants of the spike protein. Next, we describe our methods in detail and provide a list of stabilizing mutations for spike protein.

## METHODS

### Remodeling of the spike protein in different states

We utilize the crystal structure of the prefusion conformation^12^ of the spike protein (6VSB) as the template to model the spike protein through homology modeling by using MODELLER^42^. We generate 5 models and select the model with the lowest Discrete Optimized Protein Energy (DOPE)^43^, which is a statistical potential used to assess homology models in protein structure prediction, as the representative conformation of the spike protein. Since steric clashes are common in modeled and low-resolution structures, we employ Chiron^44^ to optimize the structure of the spike protein. Chiron resolves atomic clashes by performing short-DMD^38–41,45,46^ simulations on protein structures with minimal or no perturbation to the backbone. Both the template structure and the modeled structure have one RBD in up conformation and two RBDs in down conformation (1-up-2-down). We utilize PyMol^47^ to create a trimeric spike protein structure with all three RBDs in up conformation (3up) and another trimeric structure with all three RBDs in down conformation (3-down), respectively. Based on the modeled structure with all missing atoms and residues completed, we first duplicate the monomer structure with the RBD in down conformation (down-monomer) and then align it to the monomer structure with the RBD in the up conformation (up-monomer). Then, we delete the up-monomer to get the 3-down trimeric structure. Similarly, we duplicate two up-monomers in the modeled 1-up-2-down structure and then align them to the two down-monomers, respectively. Finally, we delete the two original down-monomers to get a 3-up trimeric structure of the spike protein.

### Multiple sequence alignment and Conservation score

We use ConSurf^48^ to investigate the conservation score of the spike protein. The ConSurf server is a bioinformatics tool for estimating the evolutionary conservation of amino/nucleic acid positions in a protein/DNA/RNA molecule based on the phylogenetic relations between homologous sequences.

### Molecular Dynamics Simulation

We use DMD^38–41^, an event-driven simulation which employs a discrete potential energy that relies on the calculation of atomic collisions. In DMD, the all-atom protein model interacts through the implicit solvent force-field, Medusa, which includes the van der Waals interaction, hydrogen bonds, electrostatic, dihedral and angular interactions, and those due to the effective solvent. We perform 1,000,000 step DMD simulation for the cryo-EM structure of the prefusion spike protein, with the temperature set as 0.4 using the ANDERSON thermostat. The heat exchange rate is 0.1. A cubic box (500 Å x 500 Å x 500 Å) with periodic boundary condition has been employed. For the analysis, we only consider equilibrated trajectories, discarding the first part of the trajectories that are not equilibrated due to the initial high energy. The analyses of RMSF and RMSD are performed using MDAnalysis^49^.

### Computational Mutagenesis

The spike protein structure is subject to *in silico* mutagenesis studies using Eris molecular suite. Eris’s protocol induces mutation in proteins and estimates free energies of mutant (ΔG_mut_) and wild type (ΔG_wt_) conformations. Eris^25^ performs rapid side-chain repacking and backbone relaxation around the mutation site using the Monte-Carlo algorithm and subsequently evaluates ΔG_wt_ and ΔG_mut_ using Medusa force field. Then, Eris’s algorithm computes the change in free energy of the protein upon mutation by employing the following formula: ΔΔG_mut_ = ΔG_mut_?*−*?ΔG_wt_. Finally, Eris evaluates ΔΔG_mut_ values to estimate the stabilizing (ΔΔG_mut_ < 0) or destabilizing (ΔΔG_mut_ > 0) effect of the mutations.

## RESULTS

### Protein stabilization pipeline

We propose a pipeline to automatically find stabilized mutants of proteins (Figure 1). The pipeline can be divided into two stages. In the first stage, users designate the protein of interest, and then the pipeline will analyze the 3D structure of the protein by using different metrics. In the second stage, users designate the mutation sites according to the analysis of the 3D structure of the protein, and finally the pipeline will determine the stabilizing mutations of the protein. Users can either upload the 3D structure of the protein or input the PDB ID of the protein to designate the protein of interest. In the first stage, the first step is to remodel the 3D structure of the protein of interest to complete the missing atoms and residues. We integrate MODELLER into our pipeline to remodel proteins. Next, the pipeline utilizes ConSurf^48^ to calculate the conservation score of each residue in the protein of interest. The conservation score indicates the importance of the residue in maintaining protein structure and/or function. Subsequently, the pipeline utilizes DMD to analyze the flexibility of each residue in the spike protein through RMSF. The technique has already been used to efficiently study the protein folding thermodynamics and protein oligomerization and allows for a good equilibration of the structures. Then, the pipeline will calculate SASA of residues in the protein.

**Figure 1.**
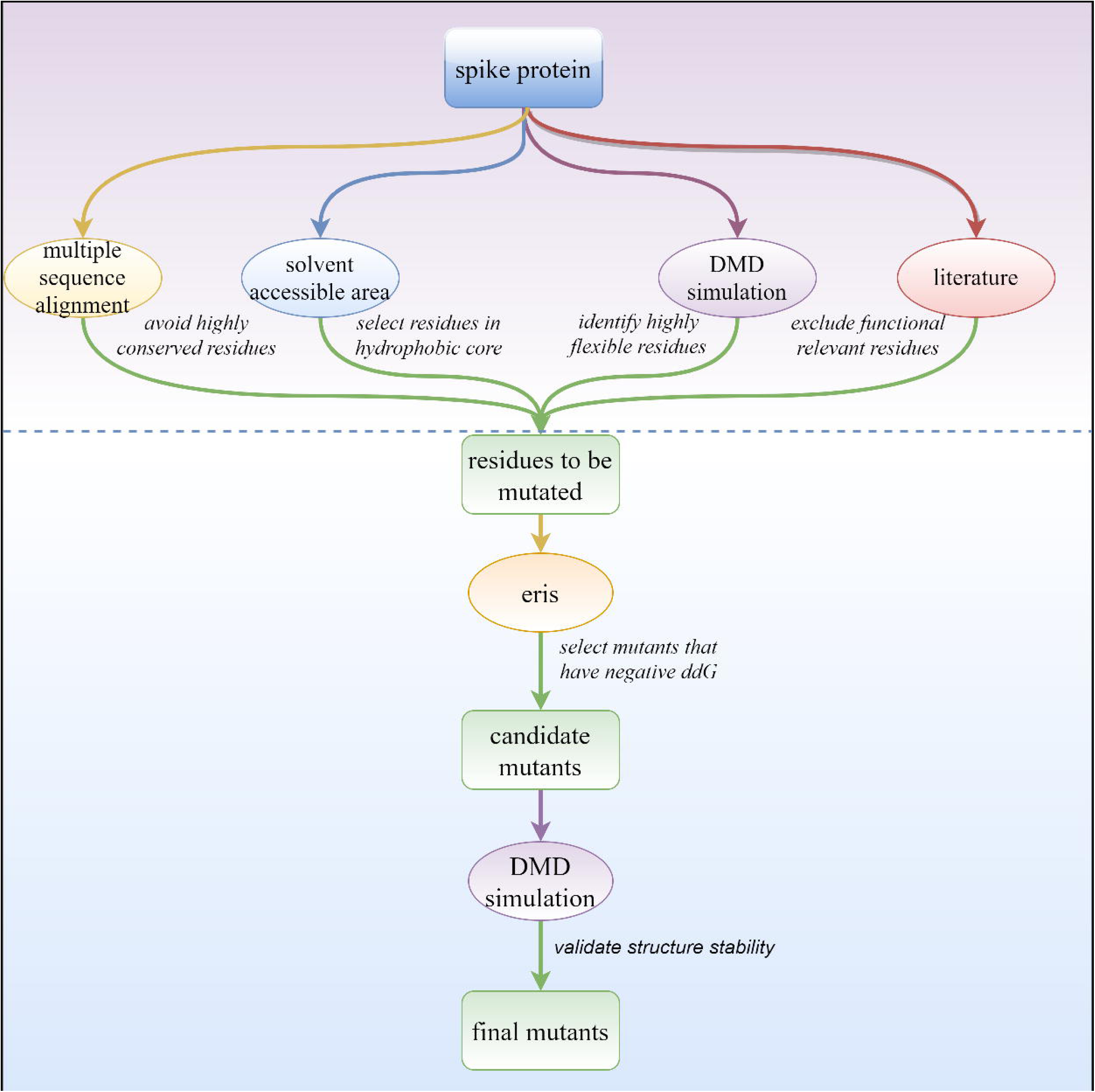
The pipeline of the stabilization of spike protein. The pipeline is roughly divided into two stages. In the first stage, users designate the protein of interest through either the 3D structure of the PDB ID. The pipeline will then analyze the conversation score, SASA, and RMSF of each residue in the protein. In the second stage, users designate the mutation sites for stabilization mutagenesis. Resides that have moderate conversation score, low SASA, and high RMSF are selected as the mutation sites. The pipeline then utilizes Eris to identify the stabilizing mutantions. Finally, the stabilization capability of these mutants is validated by DMD simulations.

In the second stage, users designate the mutation sites according to the conservation score, RMSF, and SASA. A high conservation score (≥ 7) indicates the residue may play important roles in the function or the stability of the structure of the protein; residues of high RMSF (> 3.5 Å) are likely the culprit to undermine the stability of the structure of the protein, hence we select residues that have a low conservation score or high RMSF; residues with SASA < 0.5 are considered buried and residues with SASA ≥ 0.5 are considered exposed to solvent. After the designation of the mutation sites, the pipeline utilizes Eris to determine the changes in free energies of the mutants. For each residue in the mutation sites, Eris will mutate the amino acid type of the residue to the other 19 amino acid types, iteratively. For each mutant, Eris will calculate the ΔΔG, which is the difference of the free energy of the mutant relative to the wild type. Positive ΔΔG means the mutant decreases the stability of the protein, and negative ΔΔG means the mutant can stabilize the protein. The mutants that have the lowest ΔΔG will be selected as the stabilization mutants.

### Remodeling of the spike protein structure

We utilize MODELLER^42^ to re-model the 3D structure of the spike protein by using a structure deposited to Protein DataBank (PDB), PDBID:6VSB^12^, the cryo-EM structure of the prefusion state of the spike protein, as the template structure to complete the missing atoms and residues (Figure 2A&B). Next, we use Chiron^44^, a protein energy minimization tool based on DMD, to optimize the modeled structure of the spike protein. We evaluate the modeled structure by calculating the DOPE score of each residue. DOPE of residues in the region (1147-1288) near the C-terminal are higher than that of other regions because this region is not experimentally solved in the template structure 6VSB (Figure 2C). The deposited structure 6VSB is a spike protein trimer, where one RBD is in up conformation and the other two RBDs are in down conformation. RBD domain can only bind to the ACE2 receptor when it is in the up conformation. Based on the modeled structure, we next model two spike protein structures that have all 3 RBDs in up conformation (Figure 2D) and have all 3 RBDs in down conformation (Figure 2E), respectively. All following computational mutagenesis study are performed using the structure with the 3 RBDs in down conformation.

**Figure 2.**
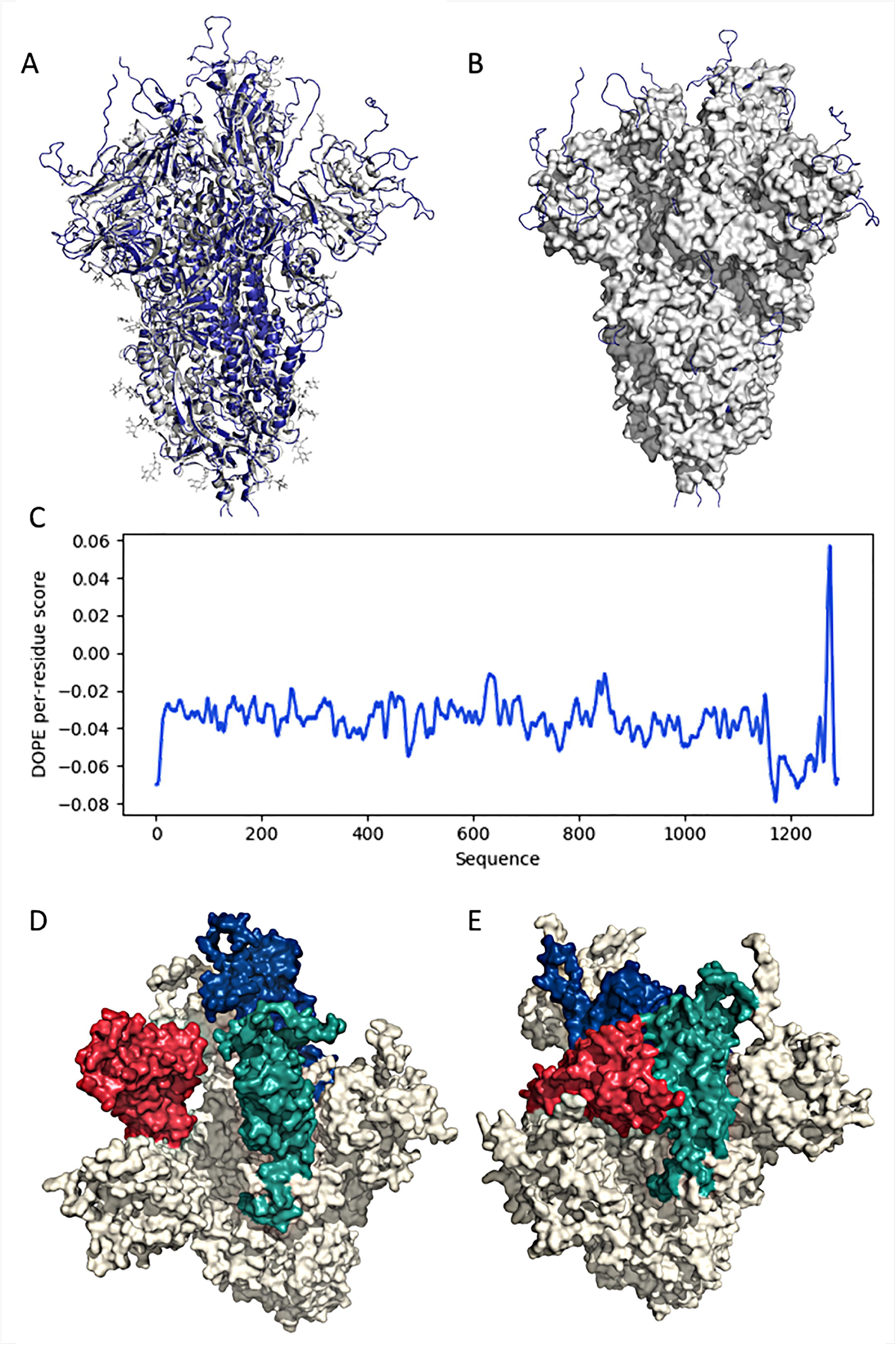
Remodeling results of the spike protein structure. (A) Comparison of the crystal structure and the remodeled structure of the spike protein. (B) The surface representation of the spike protein. The extra blue loops are the completed loops. (C) The DOPE score of each residue in the structure remodeled by MODELLER. (D) The remodeled structure of the spike protein with three RBD in up conformations. (E) The remodeled structure of the spike protein with three RBDs in down conformations.

### Validation of the protein stabilization capability of the pipeline through known mutants

To validate the ability of the pipeline to identify stabilization mutants, we use the pipeline to calculate the free energy changes of several known prefusion spike protein stabilization mutants. The 2P mutation strategy (K986P and V987P) has been proved effective for the stabilization of spike protein of SARS-COV-2 and other betacoronavirus^12,20,50^. Hsieh and coworkers^51^ have tested a large amount of mutants and found the best mutant, HexaPro, which has six beneficial prolines substitutions (F817P, A892P, A899P, A942P, K986P, and V987P) leading to ∼10-fold higher expression. We calculate the free energy change of the 2P mutant and the HexaPro mutant through Eris. The free energy change of the 2P mutant is - 6.024 kcal/mol, indicative of the more stable property of the mutant than the wild type structure. The free energy change of the HexaPro mutant is -16.143 kcal/mol, suggesting that it is even more stable than the 2P mutant. Thus, these computational calculation of the stabilities of the 2P mutant and the HexaPro mutant are in agreement with the experimental demonstration of their stability.

### Identification of new mutation sites in the spike protein

The spike protein is mainly composed of S1 and S2 subunits (Figure S1). We select residues for mutation from the NTD and RBD domains in S1, and we also select residues from the HR1 (heptad repeat 1) and CH (central helix) domains in S2. We don’t select residues from HR2 domain in S2 because the structure of the HR2 domain has not been solved in the cryo-EM structure 6VSB. At the outset, we calculate the conservation score (Figure 3A&D) of all residues by using ConSurf. Based on the conservation score, most residues in HR1/CH are conservative, while residues in NTD and RBD are prone to mutation in evolution. Next, we use Pymol^47^ to calculate SASA of all residues in the spike protein (Figure 3B&E). SASA indicates the level of residues exposed to the solvent in a protein and usually most of the functional residues are located on the protein structure’s surface^52^. All four domains have both low SASA residues and high SASA residues. Then, we perform 1,000,000 steps DMD simulation for the spike protein to calculate the RMSF of each residue (Figure 3C&F). The residues in HR1/CH have extremely low RMSF, while the residues in NTD and RBD domains have moderate to high RMSF.

**Figure 3.**
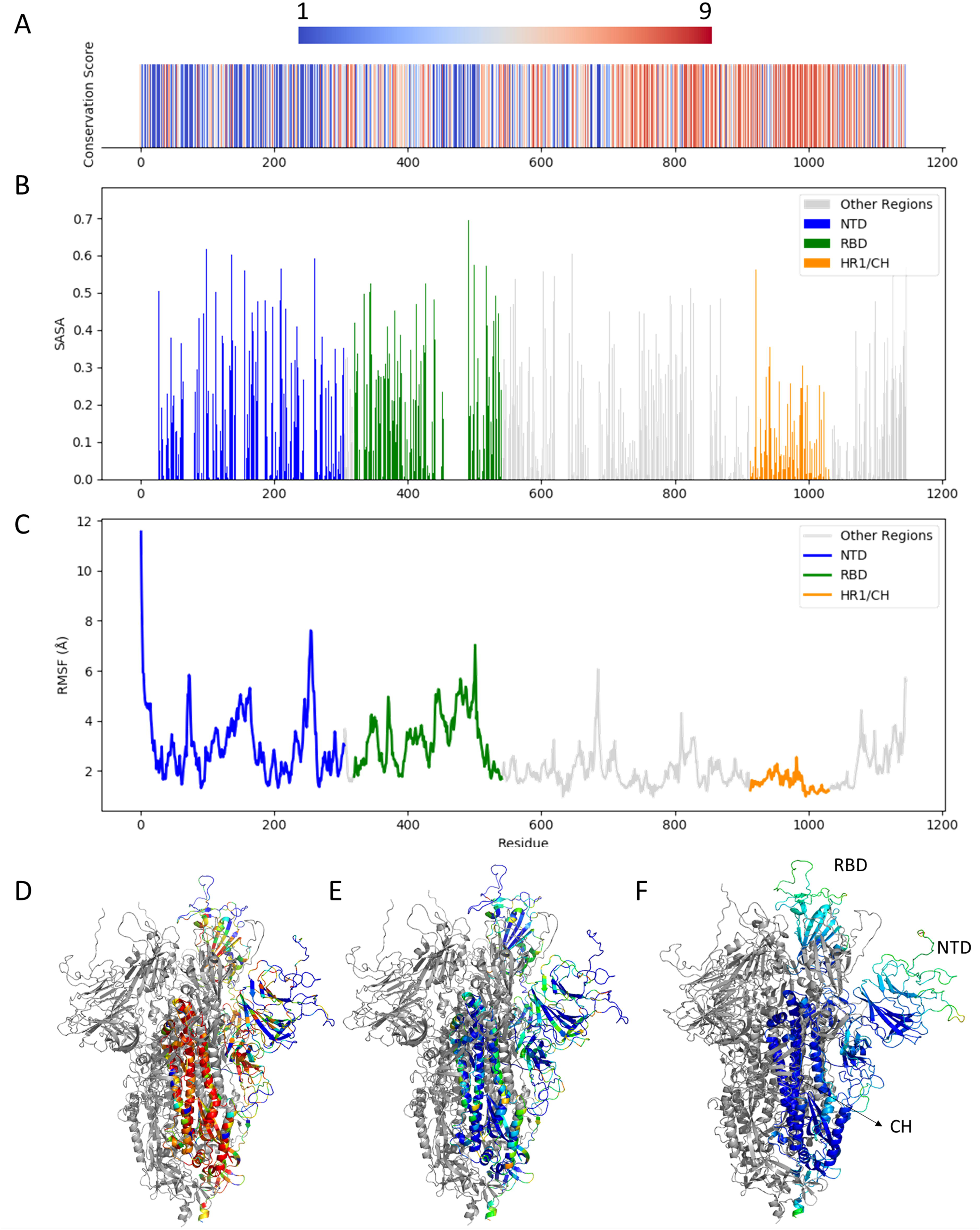
Conservation score, SASA, and RMSF of the spike protein. (A) The conservation score of residues in the spike protein. Conservation score of 9 means highly conserved, while conservation score of 1 means a highly variable position. (B) SASA of residues in the spike protein. (C) The RMSF of residues in the spike protein. (D) The 3D structures of the spike protein colored by the conservation score. The red/blue colors indicate highly conserved/highly variable residues. (E) The 3D structure of the spike protein colored SASA. The red/blue colors indicate exposed/buried residues. (F) The 3D structure of the spike protein colored by RMSF. Red means flexible and blue means frozen.

We select residues that have different conservation scores in these four domains for mutagenesis. In the NTD, we select 5 residues (Table S1) with the conservation score ranging from 1 (highly variable) to 9 (highly conservative). The SASA of these residues range from 0.01 (buried) to 0.75 (exposed). The RMSF range from 1.61 Å (frozen) to 4.88 Å (flexible). Likewise, in the other three domains (Table S2-3), we select 5 to 7 residues, respectively. These residues also have diverse conservation scores, SASA, and RMSF. Of note, to avoid affecting the function of the spike protein, these residues are all not chosen from the functional sites of the spike protein, such as the ACE2 binding site in RBD.

### Stabilization mutants of the spike protein

We utilize Eris to calculate the free energy changes of mutants relative to the wild type (Figure 4A, Figure S2&3, and Table S4-6). In the NTD, 5 residues, E169, K113, I203, R246, and L270 are selected for mutagenesis. Among them, the free energy changes of nearly all mutations of residues I203, R246, and L270 are positive, indicating that they are destabilizing the structure. In contrast, most mutations on residues E169 and K113 have negative free energy changes, suggesting that they are stabilizing the structure. The mutant that has the most negative free energy change is R246C. However, cysteine is prone to forming disulfide bond with other cysteine, which may affect the correct folding of the protein structure, so we recommend K113M to be a better choice as the stabilization mutant.

**Figure 4.**
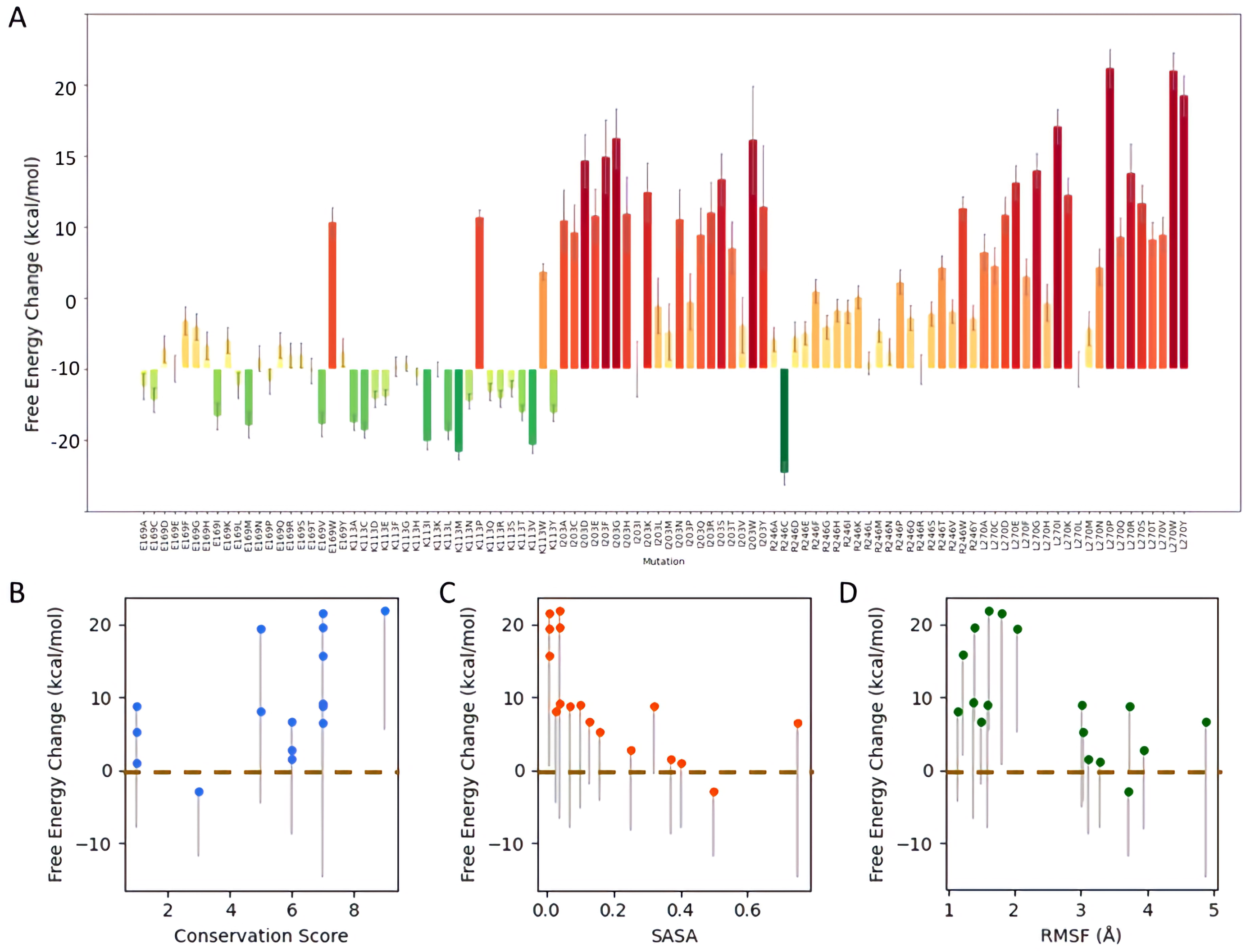
Stabilization results of the spike protein. (A) The free energy change (ΔΔG) of all mutations on the selected residues in NTD of the spike protein. (B) The correlation between free energy change and the conservation score for all mutants of the four domains. The blue dots refer to the average free energy change of all 19 mutants of each residue. The bottom end of each grey line refers to the minimum free energy change of all 19 mutants of the residue. (C) The correlation between free energy change and SASA for all mutants of residues in the four domains. (D) The correlation between free energy change and RMSF for all mutants of residues in the four domains.

In the RBD, 5 residues (A411, T415, Y505, N439, and D428) (Table S2) are selected for mutagenesis. The free energy changes calculated by Eris (Figure S2, Table S5) show that most residues have stabilization mutants, which have negative free energy changes, except for Y505. The most stable mutants are T415V and D428M.

In the HR1/CH domain, 7 residues (L948, I1018, A1026, Y1007, S1003, T961, and V976) are selected for mutagenesis as shown in Table S3. In stark contrast to NTD and RBD, most residues have extremely high free energy changes (> 30 kcal/mol), suggesting that these residues are not very good choices for mutagenesis. The high free energy changes also implicate that they may play important roles in stabilizing the structure so that they are irreplaceable to some extent. This finding is also in concert with the high conservation scores of these residues. That said, we can still find stabilization mutants for these residues, such as T961A and S1003M (Figure S3).

## DISCUSSION

Compared to experimental mutagenesis, such as random mutagenesis^53,54^ and site-directed mutagenesis^55,56^, computational mutagenesis^25^ is an efficient alternative that lays the foundation of large-scale mutation screening. However, performing computational mutation screening for all residues in the spike protein trimeric structure, which consists of 1288 residues in each monomer, is still time-consuming and inefficient, so we seek to select critical residues to perform mutagenesis. In this work, to interrogate how to select residues as mutation sites, we select residues that have different conservation scores, SASA, and RMSF. We find that the probability of stabilizing mutations for specific residues is correlated with the conservation score, SASA, and RMSF of these residues (Figure 4B-D). We calculate the average and the minimum free energy change of all 19 mutations of each residue. We find that the average free energy change of mutations of residues with either high (>5) or low (<2) conservation score are typically larger than 0. Only mutations in residues that have moderate (2∼5) conservation score have negative average free energy change, thus indicating a possibility to find stabilizing mutations. We posit that conservative residues are typically playing critical roles in structural stability or protein functioning^57^, making them a likely target for finding stabilizing mutations. However, if the conservation score of the residue is too high, the residue may be irreplaceable and any mutation will destabilize the structure. On the other hand, residues with low conservation scores may be less critical to the structural stability, reducing the chances for finding stabilizing mutations at these positions. Thus, we select residues that have moderate conservation score (2∼5) for mutagenesis. Similarly, we find that residues with large SASA or large RMSF have more stabilization mutants than residues with low SASA or low RMSF, respectively. Overall, we select residues that have moderate conservation score (2∼5), high SASA (>0.4), and high RMSF (>3.5 Å) for mutagenesis. In addition, although we only select residues in NTD, RBD, and HR1/CH domains to perform mutagenesis, residues in other regions can also be used as mutation sites. For example, the known 2P mutation strategy (K986P and V987P) has been proved effective for the stabilization of spike protein of SARS-COV-2 and other betacoronavirus^12,20,50^.

In this work, we propose a pipeline to automatically stabilize proteins through computational mutagenesis. We analyze the conservation score, RMSF, and SASA of residues in the spike protein through the pipeline. We propose criteria based on the conservation score, RMSF, and SASA to identify residues for mutation. Finally, we utilize Eris to calculate the free energy change and find stabilizing mutants.

## CODE AVAILABILITY

All source codes are deposited in: https://bitbucket.org/dokhlab/protein-stabilization.

## ACKNOWLEDGEMENTS

We acknowledge support from the National Institutes for Health 1R35 GM134864, The Huck Institutes of the Life Sciences, and the Passan Foundation. The project described was also supported by the National Center for Advancing Translational Sciences, National Institutes of Health, through Grant UL1 TR002014. The content is solely the responsibility of the authors and does not necessarily represent the official views of the NIH.

