## Supplemental information for "Prefusion spike protein stabilization through computational mutagenesis"

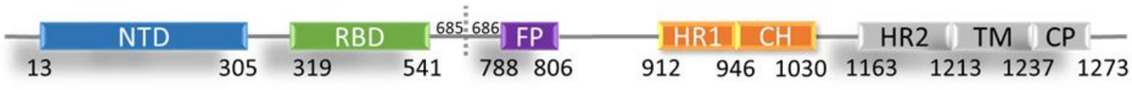


**Figure S1. The structural diagram of SARS-CoV-2's spike protein.** NTD: N-terminal domain; RBD: receptor-binding domain; FP: fusion peptide; HR1: heptad repeat 1; CH: central helix; HR2: heptad repeat 2; TM: transmembrane domain; CP: cytoplasmic; 685/686: the protease cleavage site. Structures of the domains that are colored by grey have not been solved in the prefusion spike protein cryo-EM structure 6VSB.

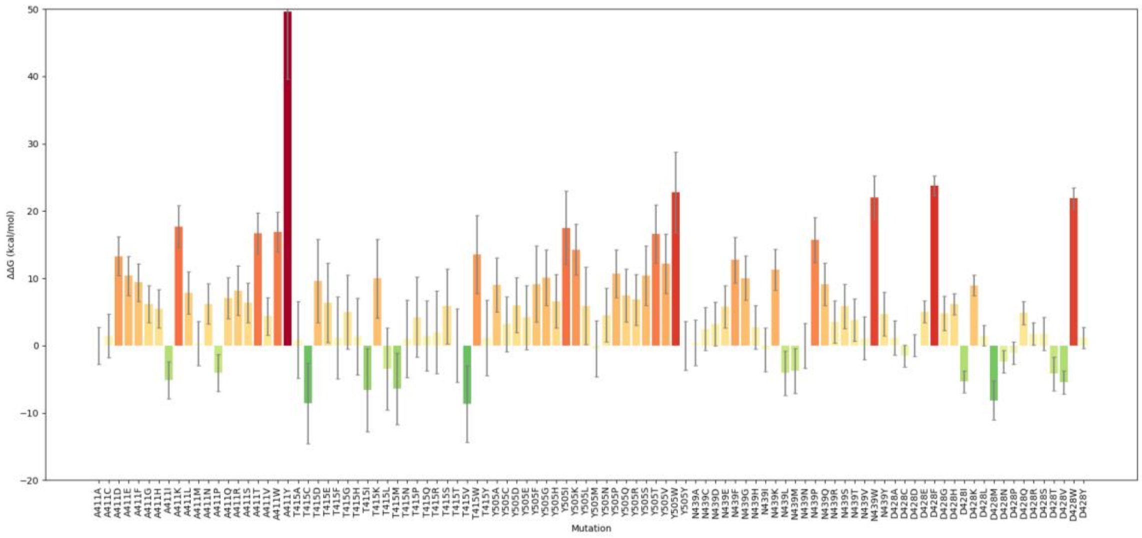


**Figure S2. Free energy changes upon mutations of selected residues in RBD domain.** The red color indicates high free energy change and the green color indicates low free energy change.

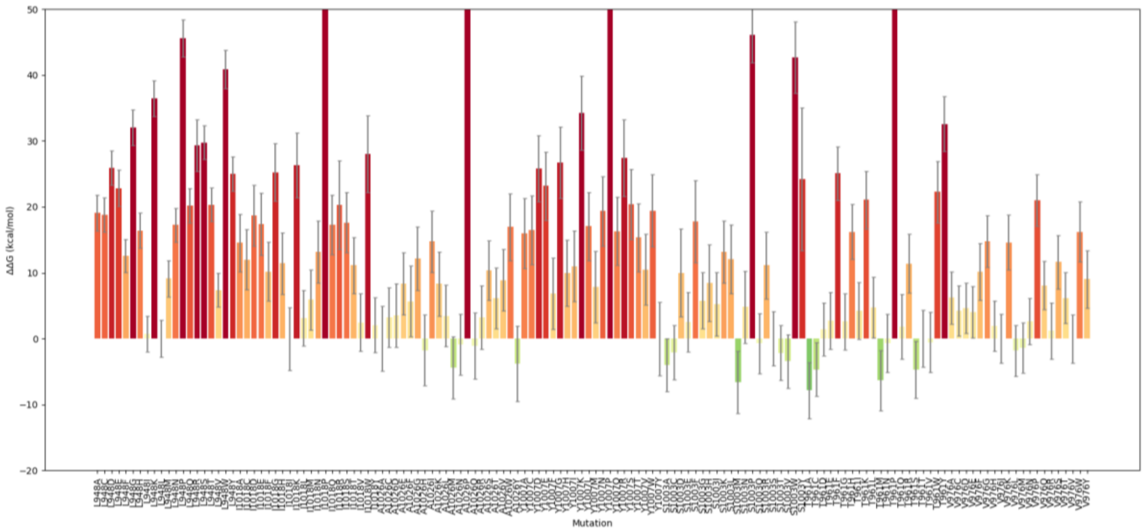


**Figure S3. Free energy changes of mutations of selected residues in HR1/CH domain.** The red color indicates high free energy change and the green color indicates low free energy change.

**Table S1. Mutation sites in NTD.**

| Residue | Conservation Score | SASA | RMSF (Å) |
|---|---|---|---|
| E169 | 1 | 0.40 | 3.29 |
| K113 | 3 | 0.50 | 3.72 |
| I203 | 5 | 0.01 | 2.04 |
| R246 | 7 | 0.75 | 4.88 |
| L270 | 9 | 0.04 | 1.61 |

**Table S2. Mutation sites in RBD.**

| Residue | Conservation Score | SASA | RMSF (Å) |
|---|---|---|---|
| A411 | 7 | 0.10 | 3.01 |
| T415 | 6 | 0.37 | 3.11 |
| Y505 | 1 | 0.32 | 3.74 |

| | | | |
|---|---|---|---|
| N439 | 1 | 0.16 | 3.04 |
| D428 | 6 | 0.25 | 3.95 |

**Table S3. Mutation sites in HR1/CH domain.**

| Residue | Conservation | SASA | RMSF (Å) |
|---|---|---|---|
| L948 | 7 | 0.01 | 1.8 |
| I1018 | 7 | 0.01 | 1.22 |
| A1026 | 5 | 0.03 | 1.15 |
| Y1007 | 7 | 0.04 | 1.39 |
| S1003 | 7 | 0.04 | 1.37 |
| T961 | 7 | 0.07 | 1.59 |
| V976 | 6 | 0.13 | 1.49 |

**Table S4. Free energy changes of the selected residues in NTD.**

| Residue | E169 | K113 | I203 | R246 | L270 |
|---|---|---|---|---|---|
| A | -2.45 | -7.51 | 20.91 | 4.23 | 16.50 |
| C | -4.38 | -8.62 | 19.26 | -14.60 | 14.57 |
| D | 2.83 | -4.18 | 29.23 | 4.58 | 21.69 |
| E | - | -3.87 | 21.46 | 5.10 | 26.23 |
| F | 6.87 | 0.41 | 29.86 | 11.01 | 12.94 |
| G | 6.00 | 0.84 | 32.42 | 5.97 | 27.82 |
| H | 3.35 | -1.01 | 21.79 | 8.34 | 9.27 |
| I | -6.66 | -10.17 | - | 8.19 | 34.12 |
| K | 4.13 | - | 24.89 | 10.18 | 24.47 |
| L | -2.32 | -8.78 | 8.83 | 0.87 | - |
| M | -7.87 | -11.69 | 5.20 | 5.46 | 5.68 |
| N | 1.53 | -4.48 | 21.13 | 2.48 | 14.30 |

| | | | | | |
|---|---|---|---|---|---|
| P | -1.67 | 21.35 | 9.43 | 12.32 | 42.27 |
| Q | 3.39 | -3.19 | 18.84 | 7.26 | 18.54 |
| R | 1.98 | -4.14 | 21.99 | - | 27.48 |
| S | 1.96 | -2.70 | 26.66 | 7.82 | 23.26 |
| T | -0.24 | -6.19 | 17.00 | 14.30 | 18.29 |
| V | -7.76 | -10.66 | 6.10 | 8.12 | 18.97 |
| W | 20.68 | 13.75 | 32.18 | 22.68 | 41.93 |
| Y | 2.40 | -6.13 | 22.75 | 7.21 | 38.44 |

**Table S5. Free energy changes of the selected residues in RBD.**

| Residue | A411 | T415 | Y505 | N439 | D428 |
|---|---|---|---|---|---|
| A | - | 0.85 | 9.03 | 0.39 | 1.17 |
| C | 1.47 | -8.54 | 3.19 | 2.46 | -1.53 |
| D | 13.28 | 9.62 | 6.03 | 3.24 | - |
| E | 10.38 | 6.39 | 4.18 | 5.78 | 5.04 |
| F | 9.38 | 1.14 | 9.15 | 12.74 | 23.76 |
| G | 6.14 | 4.98 | 10.09 | 10.05 | 4.82 |
| H | 5.48 | 1.37 | 6.60 | 2.73 | 6.21 |
| I | -5.11 | -6.63 | 17.50 | -0.59 | -5.34 |
| K | 17.72 | 9.98 | 14.25 | 11.25 | 8.98 |
| L | 7.85 | -3.46 | 5.92 | -4.09 | 1.49 |
| M | 0.32 | -6.42 | -0.43 | -3.76 | -8.12 |
| N | 6.19 | 1.00 | 4.52 | - | -2.38 |
| P | -4.08 | 4.25 | 10.73 | 15.67 | -1.10 |
| Q | 7.04 | 1.44 | 7.44 | 9.14 | 4.85 |
| R | 8.16 | 1.99 | 6.83 | 3.49 | 1.75 |
| S | 6.35 | 5.84 | 10.39 | 5.86 | 1.79 |
| T | 16.70 | - | 16.59 | 3.79 | -4.18 |
| V | 4.39 | -8.68 | 12.17 | 1.11 | -5.44 |

| | | | | | |
|---|---|---|---|---|---|
| W | 16.86 | 13.54 | 22.78 | 21.97 | 21.87 |
| Y | 49.63 | 1.16 | - | 4.71 | 1.13 |

**Table S6. Free energy changes of the selected residues in HR1/CH domain.**

| Residue | L948 | A1026 | Y1007 | S1003 | T961 | V976 |
|---|---|---|---|---|---|---|
| A | 19.11 | 14.56 | - | 15.97 | -4.01 | -7.83 |
| C | 18.79 | 12.01 | 3.23 | 16.50 | -2.10 | -4.66 |
| D | 25.96 | 18.67 | 3.51 | 25.80 | 9.98 | 1.43 |
| E | 22.86 | 17.40 | 8.37 | 23.20 | 2.51 | 2.73 |
| F | 12.61 | 10.20 | 5.68 | 6.83 | 17.78 | 25.10 |
| G | 32.05 | 25.23 | 12.21 | 26.68 | 5.78 | 2.58 |
| H | 16.41 | 11.42 | -1.75 | 9.95 | 8.47 | 16.19 |
| I | 0.73 | - | 14.75 | 11.00 | 5.22 | 4.25 |
| K | 36.48 | 26.28 | 8.31 | 34.25 | 13.18 | 21.08 |
| L | - | 3.14 | 3.48 | 17.05 | 12.09 | 4.71 |
| M | 9.13 | 5.92 | -4.36 | 7.88 | -6.60 | -6.34 |
| N | 17.26 | 13.17 | -0.87 | 19.35 | 4.83 | -0.68 |
| P | 45.60 | 58.93 | 68.02 | 66.87 | 46.08 | 56.30 |
| Q | 20.20 | 17.31 | -1.07 | 16.30 | -0.71 | 1.82 |
| R | 29.35 | 20.30 | 3.25 | 27.44 | 11.12 | 11.39 |
| S | 29.78 | 17.60 | 10.32 | 20.39 | - | -4.70 |
| T | 20.33 | 11.13 | 6.11 | 15.34 | -2.15 | - |
| V | 7.39 | 2.47 | 8.88 | 10.51 | -3.40 | -0.54 |
| W | 40.90 | 28.05 | 16.97 | 19.43 | 42.71 | 22.26 |
| Y | 24.99 | 2.05 | -3.78 | - | 24.19 | 32.58 |
